# Microtubules regulate pancreatic beta cell heterogeneity via spatiotemporal control of insulin secretion hot spots

**DOI:** 10.1101/2020.06.12.148668

**Authors:** Kathryn P. Trogden, Hudson McKinney, Xiaodong Zhu, Goker Arpag, Thomas G. Folland, Anna B. Osipovich, Mark A Magnuson, Marija Zanic, Guoqiang Gu, William R. Holmes, Irina Kaverina

**Author notes:** Corresponding author: Irina Kaverina.

## Abstract

Heterogeneity of glucose-stimulated insulin secretion (GSIS) in pancreatic islets is physiologically important but poorly understood. Here, we utilize whole mouse islets to determine how microtubules affect secretion toward the vascular extracellular matrix. Our data indicate that microtubule stability in the β-cell population is heterogenous, and that cells with more stable microtubules secrete less in response to a stimulus. Consistently, microtubule hyper-stabilization prevents, and microtubule depolymerization promotes β-cell activation. Analysis of spatiotemporal patterns of secretion events shows that microtubule depolymerization activates otherwise dormant β-cells via initiation of secretion clusters (hot spots). Microtubule depolymerization also enhances secretion from individual cells, introducing both additional clusters and scattered events. Interestingly, without microtubules, the timing of clustered secretion is dysregulated, extending the first phase of GSIS. Our findings uncover a novel microtubule function in tuning insulin secretion hot spots, which leads to accurately measured and timed response to glucose stimuli and promotes functional β-cell heterogeneity.

## Introduction

Insulin secretion in pancreatic β-cells is tightly regulated and highly heterogeneous. The major stimulator for secretion is glucose, with other nutritional and neuronal signals modulating the response. The prevailing view is that glucose influx results in increased glucose metabolism, higher levels of the cytoplasmic ATP/ADP ratio, and increased levels of metabolite intermediates. The increased ATP/ADP ratio results in the closure of K_ATP_ channels, β-cell depolarization, and Ca^2+^ influx that triggers vesicular-cell membrane fusion. Metabolite intermediates promote secretion via known (e.g. microtubule dependent vesicular biogenesis, movement, and docking) and unknown mechanisms. Intriguingly, β-cells secrete on the order of tens of granules per cell in response to each round of high-glucose stimulus, which last for hours, despite having thousands of available granules (Dean, 1973, Olofsson et al., 2002, Rorsman and Renstrom, 2003). Glucose-stimulated insulin secretion (GSIS) has a characteristic bi-phasic kinetics with a rapid, high peak within minutes after the stimulation (first phase) followed by a sharp decrease and a slow, lower secreting second phase. The mechanisms underlying this timing are believed to be associated with the availability of secretion-ready insulin vesicles that do not depend on changes in Ca^2+^ levels (Gaisano, 2017). m

It has long been known that β-cells in islets are functionally heterogeneous, so that different subpopulations of β-cells have variable insulin secretion capacities and timing in their responses to high glucose stimuli (Salomon and Meda, 1986, Van Schravendijk et al., 1992, Pipeleers, 1992, Bosco et al., 1995, Wojtusciszyn et al., 2008, Bonner-Weir and Aguayo-Mazzucato, 2016, Gutierrez et al., 2017, Pipeleers et al., 2017). This heterogeneity can be explained in part by differences in the Ca^2+^ influx rate, because the response starts from a regulated Ca^2+^ influx in a small number of “hub cells” and is transmitted further through the involvement of gap junctions (Johnston et al., 2016). However, the secretion capacity of β-cells remains heterogeneous even when the glucose-induced Ca wave becomes relatively uniform (Benninger and Hodson, 2018) or when uniform Ca^2+^ influx is enforced by KCl (Li et al., 2011). Differential sensitivity to glucose cannot completely explain this observation, because approximately 25% β-cells remain unresponsive regardless of the glucose concentration (Hoang Do and Thorn, 2015). In addition, gene expression patterns and proliferation capacity are also heterogeneous in the β-cell population (Dorrell et al., 2016, Bader et al., 2016). It has been proposed that β-cell heterogeneity can result from differences in age, disease state, β-cell maturity, and location within the islet (Pipeleers et al., 2017, Gutierrez et al., 2017, Aguayo-Mazzucato et al., 2017). Specific locations that might influence secretion activity of individual cells include their attribution to a specific islet cell layer (Ballian and Brunicardi, 2007, Stefan et al., 1987, Dean and Matthews, 1970), their position in relation to other islet cell subtypes (Wojtusciszyn et al., 2008, Pipeleers et al., 1982, van der Meulen et al., 2015, Efendic and Luft, 1975) and to vasculature (Ballian and Brunicardi, 2007, Low et al., 2014).

In additon to secretory heterogeneity in the β-cell population, uneven distribution of secretion was also observed at subcellular levels. Designation of insulin secretion to specialized loci at the cell membrane was shown to be essential for the pathophysiology of type 2 diabetes (Fu et al., 2019). The sites of preferential insulin secretion depend on the cellular location of L-type voltage-dependent calcium channels in combination with molecular tethers and other exocytotic proteins (Ohara-Imaizumi et al., 2019b, Bokvist et al., 1995). This molecular machinery resembles the composition of “active zones” or areas of high exocytosis in neurons (Garner et al., 2000) and is thought to underlie the hot spots of secretion at the plasma membrane (Landis et al., 1988). Interestingly, some major components of the hot spot machinery (e.g. ELKS) are assembled at the membrane in response to integrin activation by vascular extracellular matrix (ECM) proteins such as laminin (Nishimune, 2012, Hotta et al., 2010), and in islets they are preferentially found at sites of β-cell contact with the vasculature (Ohara-Imaizumi et al., 2005, Low et al., 2014). However, as most β-cells in an in-situ islet have discrete points of contact with the capillaries (Low et al., 2014), it is clear that positioning and ECM-dependent activation of secretion hot spots is insufficient for the differences in secretion activity of individual cells. Thus, the cellular mechanisms that drive the functional β-cell heterogeneity are still not well understood.

Recently, we have shown that the secretion capacity of β-cells is regulated by the microtubule (MT) cytoskeleton, which is uniquely structured to help tune β-cell function (Zhu et al., 2015). Unlike in many other cell types, MTs in β-cells do not radiate from a central point in the cell, and instead form a dense mesh-like network (Zhu et al., 2015, Trogden et al., 2019, Bracey et al., 2020). This network configuration is, to a large extent, due to the fact that most MTs in β-cells originate at the Golgi apparatus (Zhu et al., 2015, Trogden et al., 2019). The large surface area of the Golgi acts as a MT organizing center (MTOC), leading to an unusual non-radial MT network appearance. Intriguingly, optimal β-cell function depends on a finely tuned balance of MT assembly and disassembly, which can both promote and restrict secretion capacity of a cell. On the one hand, Golgi-derived MTs function aids the budding of new insulin granules from the Golgi (Trogden et al., 2019). High glucose stimuli lead to an increase in Golgi-derived MT nucleation, which is necessary to replenish insulin granule content after a secretion pulse (Zhu et al., 2015, Trogden et al., 2019). Long-term loss of this MT subpopulation causes degranulation of the β-cell as vesicle budding is less efficient (Trogden et al., 2019). Thus, glucose-dependent MT nucleation is a critical factor supporting the capacity of β-cells to secrete. On the other hand, MTs act to prevent over-secretion in functional β-cells, which contain excessive amounts of insulin granules(Zhu et al., 2015). This function relies on the configuration of MT networks in β-cells, where the non-radial mesh in the cell interior prevents directional granule movement, and the extremely stable MT bundles extending along the plasma membrane serve as tracks for granule withdrawal from docking sites (Bracey et al., 2020). Upon glucose stimulation, these pre-existing MTs are destabilized and partially depolymerized (Ho et al, 2020, in press), allowing for release of a subset of granules. This regulated dynamicity of the MT network is vital for the dosage of GSIS at each stimulus: loss of all MTs acutely leads to over-secretion, while hyper-stabilization of MTs greatly suppresses it (Zhu et al., 2015, Lacy et al., 1972, Howell et al., 1982, Hill and Rhoten, 1983). These effects are only seen after GSIS stimulation but not at basal glucose conditions, indicating the leading role for other mechanisms in secretion triggering (Zhu et al., 2015, Trogden et al., 2019). In functional β-cells which are able to secrete in response to a glucose stimulus, the effects of MT destabilization are finely tuned because simultaneous increase in MT nucleation promptly replaces depolymerizing MTs (Zhu et al., 2015). The intriguing combination of positive (long-term) and negative (short-term) MT regulation of secretion may be responsible for the early controversy on the role of MTs in insulin secretion (Lacy et al., 1968, Lacy et al., 1972, Malaisse et al., 1974, Howell et al., 1982, Hill and Rhoten, 1983, Mourad et al., 2011). It also makes the MT cytoskeleton a candidate for differential regulation of β-cells activity underlying functional heterogeneity.

In this paper we investigate the role of the MT cytoskeleton in β-cells as a factor in the differential secretory response of β-cells to glucose stimulation. Our computational analysis of insulin secretion at the single-cell level toward the vascular ECM in whole islets shows that this regulation occurs via secretion hot spot activity. Our findings indicate that the presence of stable MTs attenuates initiation of secretion hot spots in both otherwise dormant and already active β-cells, thus contributing to functional heterogeneity of insulin secretion. We also show that MTs regulate the timing of insulin secretion by restricting hot spot activity to the first phase of GSIS.

## Results

### Microtubule stability in pancreatic islet β-cells is heterogenous

We have previously found that MTs serve as a critical regulator of both phases of β-cell glucose response and insulin secretion (Zhu et al., 2015, Trogden et al., 2019, Ho et al, 2020, in press). To test whether MT regulation plays a role in the functional heterogeneity of β-cell population, we analyzed MT stability in mildly disseminated mouse islet β-cells plated on vascular ECM by three independent approaches.

First, MT stability was evaluated utilizing immunostaining for Glu-tubulin (also known as detyrosinated tubulin), a marker of long-lived or stable MTs (Figure 1 A-F, β-cells outlined in yellow). β-cells were identified by expression of nuclear-localized mApple marker (see materials and methods). Consistent with our previous findings, Glu-tubulin content was high in β-cells at basal glucose conditions, but decreased following a high glucose stimulus (Zhu et al., 2015, Ho et al, 2020, in press, Figure 1G, H). There was no detectable difference in overall polymerized tubulin content upon glucose stimulus (Figure 1- figure supplement 1A), likely due to glucose-stimulated nucleation of new MTs (Trogden et al., 2019), which replace destabilized MTs (Zhu et al., 2015, Trogden et al., 2019). Accordingly, the ratio of Glu-tubulin to tubulin per cell, which reflects the proportion of stable MTs within the MT network, decreased after high glucose stimulation (Figure 1J, K).

**Figure 1.**
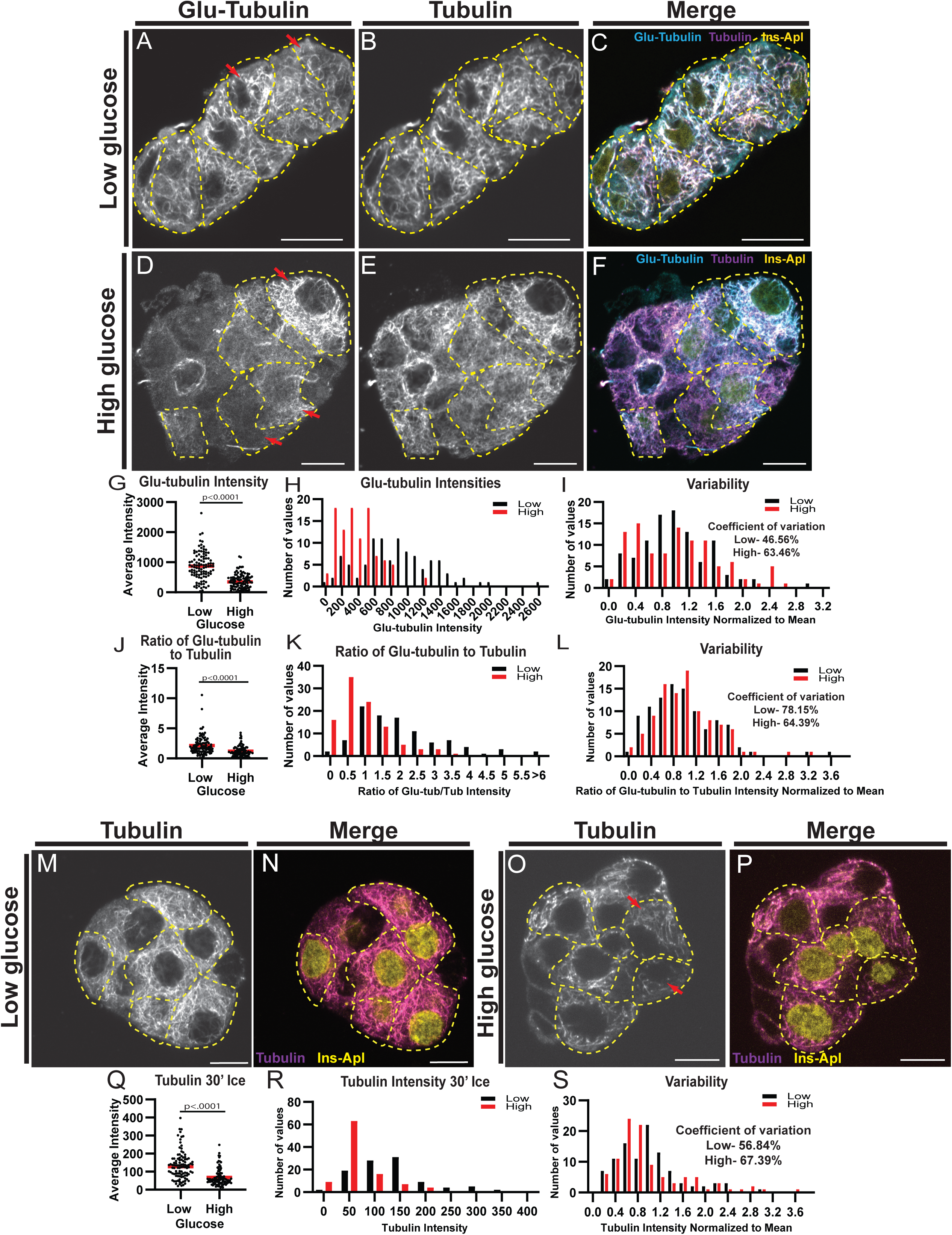
MT stability decreases in high glucose but remains heterogenous. A-F) Disseminated islets treated with low (A-C) and high (D-F) glucose stained for Glu-tubulin (A,D) and tubulin (B,E). β-cells (dashed yellow line) were identified using Ins-Apl red nuclei (yellow, C,F). Red arrows point to differences between cells. Merge (C,F) shows Glu-tubulin (cyan), tubulin (magenta) and red nuclear expression of Ins-Apl (yellow). Single slice from the bottom of the cells. Scale bars 10 µm. G) Scatterplot of Glu-tubulin average intensity for each cell. Mean, red bar. Students’ t-test, p<0.0001. n=101 cells per condition. H) Histogram of Glu-tubulin average intensity in low (black) and high (red) glucose. Bin=100. n=101 cells per condition. I) Histogram of Glu-tubulin average intensity normalized to the mean of each low (black) and high (glucose). Bin=0.2. Coefficient of variation= standard deviation/mean. n=101 cells per condition. J) Scatterplot of Glu-tubulin to tubulin ratio of average intensity for each cell. Mean, red bar. Students’ t-test, p<0.0001. n=101 cells per condition. K) Histogram of Glu-tubulin to tubulin ratio of average intensity in low (black) and high (red) glucose. Bin=0.5, overflow bin of >6. n=101 cells per condition. L) Histogram of Glu-tubulin to tubulin ratio of average intensity normalized to the mean of each low (black) and high (glucose). Bin=0.2. Coefficient of variation= standard deviation/mean. n=101 cells per condition. M-P) Disseminated islets placed on ice for 30 minutes in low (M-N) and high glucose (O-P) and stained for tubulin (M,O). β-cells (dashed yellow line) were identified using red nuclear expression of Ins-Apl (yellow, N,P), merged with tubulin (magenta N,P). Red arrows point to differences between cells. Single slice from the bottom of the cells. Scale bar 10 µm. Q) Scatterplot of tubulin average intensity for each cell after 30 minutes in high glucose. Mean, red bar. Students’ t-test, p<0.0001. n=100-101 cells per condition R) Histogram of tubulin average intensity in low (black) and high (red) glucose after 30 min on ice. Bin=50. n=100-101 cells per condition S) Histogram of tubulin average intensity normalized to the mean of each low (black) and high (glucose) after 30 min on ice. Bin=0.2. Coefficient of variation= standard deviation/mean. n=100-101 cells per condition.

Interestingly, the density of the MT cytoskeleton is heterogeneous within the β-cell population. While not changed by glucose stimulation, the overall tubulin polymer content was variable in the β-cell population as indicated by the intensity distributions (Figure 1- figure supplement 1B and C) and coefficients of variation (58.72% in low glucose and 60.42% in high glucose). MT stability was also highly variable, as evident from the Glu-tubulin staining, which ranged from barely detectable levels to highlighting essentially the whole MT cytoskeleton (Figure 1A, D, compare cells with red arrows). While the amount of Glu-tubulin decreased after glucose stimulation, shifting the histogram of Glu-tubulin intensities per cell to the left (Figure 1H), the degree of variation amongst the beta cells was retained as indicated by the distributions of Glu-tubulin intensity normalized to the mean for each condition (Figure 1I), and by high coefficients of variation (48.56% in low glucose and 63.46% in high glucose). The proportion of stable MTs within the MT network (Glu-tubulin to tubulin) also remained variable in high glucose (Figure 1 K, L; coefficient of variation: 78.15% in low glucose and 64.39% in high glucose).

As a second measure of MT stability in β-cells, we subjected the cells to ice treatment for 30 minutes. Since MTs are temperature-sensitive and only stable MTs will remain after ice treatment, this well-established assay is used to directly assess MT stability. After extraction of free tubulin, fixation and immunostaining, the tubulin content per cell was measured to evaluate the amount of cold-resistant MTs remaining (Figure 1M-P). In basal glucose conditions, many cells contained MTs stable enough to be retained after ice treatment (Figure 1M, N). Following high glucose stimulation, the amount of cold-stable MTs significantly decreased (Figure 1Q, R), consistent with our previous findings and data described above (Zhu et al., 2015, Ho et al, 2020, in press). Most of the remaining MTs were positive for Glu-tubulin, as expected for long-lived stable MTs (Figure 1- figure supplement 1D-K). Importantly, the variation in cold-resistant MTs within the β-cell population was high both in low and high glucose (Figure 1S), confirming that MT stability is highly variable in β-cell population.

As a third test of MT stability, we utilized a method where live cells were permeabilized prior to fixation to release all free tubulin, which acutely decreases tubulin concentration in the cell (Khawaja et al., 1988). This prevents tubulin polymerization so that dynamic MTs are lost through depolymerization without being replaced, leaving only stable MTs in a cell (Khawaja et al., 1988). Immunofluorescent detection of MTs after this treatment resulted in a similar variation in MTs amounts retained in cells and Glu-tubulin staining as the cold treatment (Figure 1- figure supplement 1L-N).

These three assays indicated that, similar to insulin secretion, MT stability is heterogeneous between different β-cells. While some cells have highly stable MT networks, neighboring cells can have less stable networks. This phenomenon is even more obvious after high glucose stimulation when MT stability is decreased in the entire population, but remains highly variable.

### MT stability affects insulin secretion heterogeneity

To assess whether MT regulation affects the functional heterogeneity of β-cells, we next analyzed insulin secretion from distinct cells within the β-cell population. To detect single-cell insulin secretion in real time we use a cell impermeant zinc dye that becomes fluorescent upon zinc binding, and at each exocytic event, highlights the release of zinc co-packaged with insulin in granules (Zhu et al., 2015, Gee et al., 2002). Total internal reflection fluorescent (TIRF) microscopy was used to analyze secretion in flattened intact islets attached to a vascular ECM to imitate secretion towards vasculature (See materials and methods, Figure 2- figure supplement 1A, (Gan et al., 2018, Low et al., 2014, Bonner-Weir, 1988). Individual secretion events appear as flashes of bright fluorescence, the intensity of this fluorescence over time resembles a Gaussian curve with only background signal, then an initially tight circle of signal that dissipates after vesicle exocytosis (Figure 2A, A’). Analysis started within 2 minutes after glucose stimulation to register active GSIS. For quantification, image sequences were processed for better signal/noise ratio, β-cells were identified by the presence of red fluorescence in the nuclei, and cell outlines were detected by bright-field imaging.

**Figure 2.**
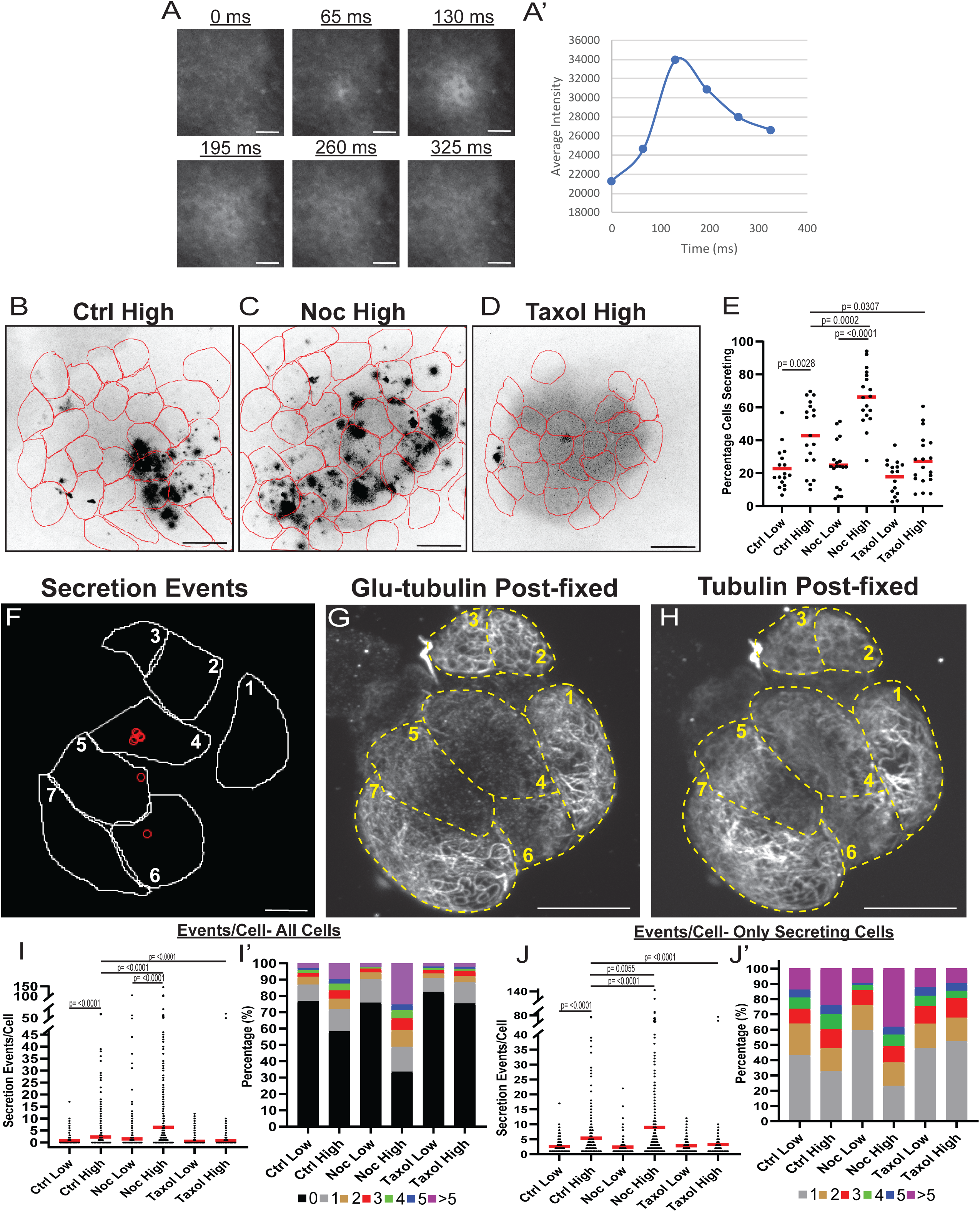
MT instability increases β-cell activation and insulin secretion. A-C) Time projections of islets from Video 1-3 (high glucose) inverted. Fluozin-3 flashes are represented as black areas. Cell borders identified via pre-assay imaging (see materials and methods) overlaid in red. Islets were pre-incubated in DMSO (control, A), nocodazole (B) or taxol (C) and stimulated with 20mM glucose. Scale bars, 100 µm. D) Example images of a single secretion event pre-processing. Secretion signal starts at 65ms and dissipates out. D’) Graph of the average intensities of a circular ROI from images in panel D. E) Graph of the percentage of cells in each field of view with at least one secretion event. Red bars, mean. One-way ANOVA and multiple comparison tests, p value as indicated. N= 16-19 islets. F) Cell outlines (white line) and secretion events (Fluozin-3 flashes, red circles) from a disseminated islet in 20 mM glucose. Scale bar 10 µm. G-H) Disseminated islet from E post-fixed following TIRF imaging for Glu-tubulin (F) and tubulin (G). Cells (yellow dashed lines) correspond to cells in E with the same number. Single slice from the bottom of the cells. Scale bar 10 µm. I) Graph of secretion events per cell in the field of view, with all cells whether activated during the movie or not. Red bar, mean. Kruskal-Wallis test non-parametric and multiple comparison tests, p value as indicated, N= 495-637 cells from 16-19 islets. I’) Cells from panel H, graphed as a stacked histogram of the percentage of total cells per condition that had each number of secretion events. J) Graph of secretion events per cell only including cells with at least one event during the duration of the movie. Red bar, mean. Kruskal-Wallis test non-parametric and multiple comparison tests, p value as indicated, N= 88-407 cells from 16-19 islets. J’) Cells from panel I, graphed as a stacked histogram of the percentage of cells that had each number of secretion events

To address heterogeneity of β-cell activity in our experimental setup, we first measured the percentage of secreting β-cells. In control islets treated with high glucose, the number of active cells increased as expected (Figure 2B, E, Figure 2- figure supplement 1B and Video 1 (Low et al., 2013). However, a subset of β-cells remained inactive even after stimulation (Figure 2E, 60% of cells), resembling the fraction of cells with a high content of stable MTs (59% of cells with >300 average intensity of glu-tubulin, Figure 1).

To assess the role of MTs in the distribution of secretion activity in the β-cell population, we pre-incubated islets in the presence of either the MT depolymerizing drug nocodazole, which completely eliminates cellular MTs (Figure 2- figure supplement 1I), or the MT-stabilizing drug taxol, which hyper-stabilizes MT networks in cells (Figure 2- figure supplement 1J). As we have previously observed (Zhu et al., 2015), nocodazole increased insulin secretion specifically in high glucose-stimulated islets (Figure 2C, E, Figure 2- figure supplement 1C and Video 2). Strikingly, we also observed that a larger sub-population of β-cells (on average, 66% of cells) was activated, compared to only 42% in control islets (Figure 2E, compare Figure 2B and C). Addition of taxol also affected insulin secretion as we have previously observed (Zhu et al., 2015), blunting it in high glucose (Figure 2D, E, Figure 2- figure supplement 1D, Video 3). The percent of secreting cells in taxol was only 27%, similar to the percentage observed in low glucose (Figure 2D, E, Figure 2- figure supplement 1D), where MTs are intrinsically stable (Figure 1).

These results indicate that the modulation of MT stability can change the insulin secretion activity of β-cells within the population. Since in control islets cells have variable MT stability levels (Figure 1), we explored a potential connection between MT stability in individual cells and their ability to secrete. This connection was assessed by correlative microscopy of the FluoZin3-detected secretion pattern with post-assay Glu-tubulin immunostaining in lightly disseminated islets. Interestingly, secretion was not detected in cells with a high content of Glu-tubulin (Figure 2F-H and Figure 2- figure supplement 1E-G, see materials and methods), indicating a direct correlation between MT dynamicity and ability of β-cells to secrete in high glucose.

We next tested whether MTs cause differences between active β-cells by assessing how modulation of MTs influences the number of secretion events per cell. We found that high glucose caused an increase in the number of events per cell in both control and nocodazole-treated islets, but not taxol-treated islets (Figure 2I, I’, J, J’ (Low et al., 2013)). Importantly, the number of events per cell and the fraction of highly-secreting cells in high glucose were significantly increased by nocodazole as compared to control, suggesting that MT presence attenuates glucose-stimulated secretion in individual cells (Figure 2I, I’, J, J’). Consistent with this result, under all conditions of high MT stability (low glucose in all conditions and high glucose in taxol) secretion activity is distributed similarly throughout the cell population (Figure 2I’, J’). Taken together, these data indicate that insulin secretion capacity on both the cellular and population levels is decreased by MT presence, and even more so, by MT stability. Furthermore, in terms of β-cell activation, MTs provide a mechanism that supports β-cell heterogeneity. While normally the glucose-stimulated β-cell population is highly heterogeneous, changing MT stability in either direction leads to a more homogenous population with either most cells secreting (nocodazole, no MTs) or most cells not secreting (taxol, hyper-stabilized MTs). It is also evident that modulation of MT dynamics decreases but does not completely eliminate variability in β-cell activity, which is in an agreement with the existence of other, MT-independent, mechanisms of heterogeneity (reviewed in (Gutierrez et al., 2017, Pipeleers et al., 2017).

### MT stability regulates β-cell activation by suppressing insulin secretion “hot spots”

To better understand MT-dependent regulation of secretion in individual cells versus a cell population, we analyzed their spatial and temporal secretion patterns. In agreement with the existing evidence that insulin secretion occurs in hot spots, defined by patches of VDCCs and other active zone proteins on the membrane (Yuan et al., 2015b, Ohara-Imaizumi et al., 2019b, Ohara-Imaizumi et al., 2005, Gandasi et al., 2017), we observed a distinct distribution of secretion events to certain preferred areas of a cell (Figure 2B-D, Videos 1-3). Analyzing the nearest neighbor distances between secretion events within cells, we detected a bias towards smaller distances, with over 50% of events in all conditions occurring within 1.5 µm of each other (Figure 3A). We then performed a density-based scanning algorithm to identify clustering, defined as secretion events occurring within 1.5 µm diameter areas (Figure 3B-D, Figure 3- figure supplement 1B-D). To determine the expected frequency of clustered events occurring by chance (i.e. multiple unrelated events occurring close to each other by coincidence), we computationally simulated random secretion events in *in silico* cells. Results (Figure 3- figure supplement 1A) show that clusters of two events within 1.5 µm of each other would be relatively prevalent (one or more per cell) with just 10 randomly distributed secretion events, while larger clusters would be uncommon (less than one per cell). Thus, only cluster sizes of three and more events were included in the data presented below.

**Figure 3.**
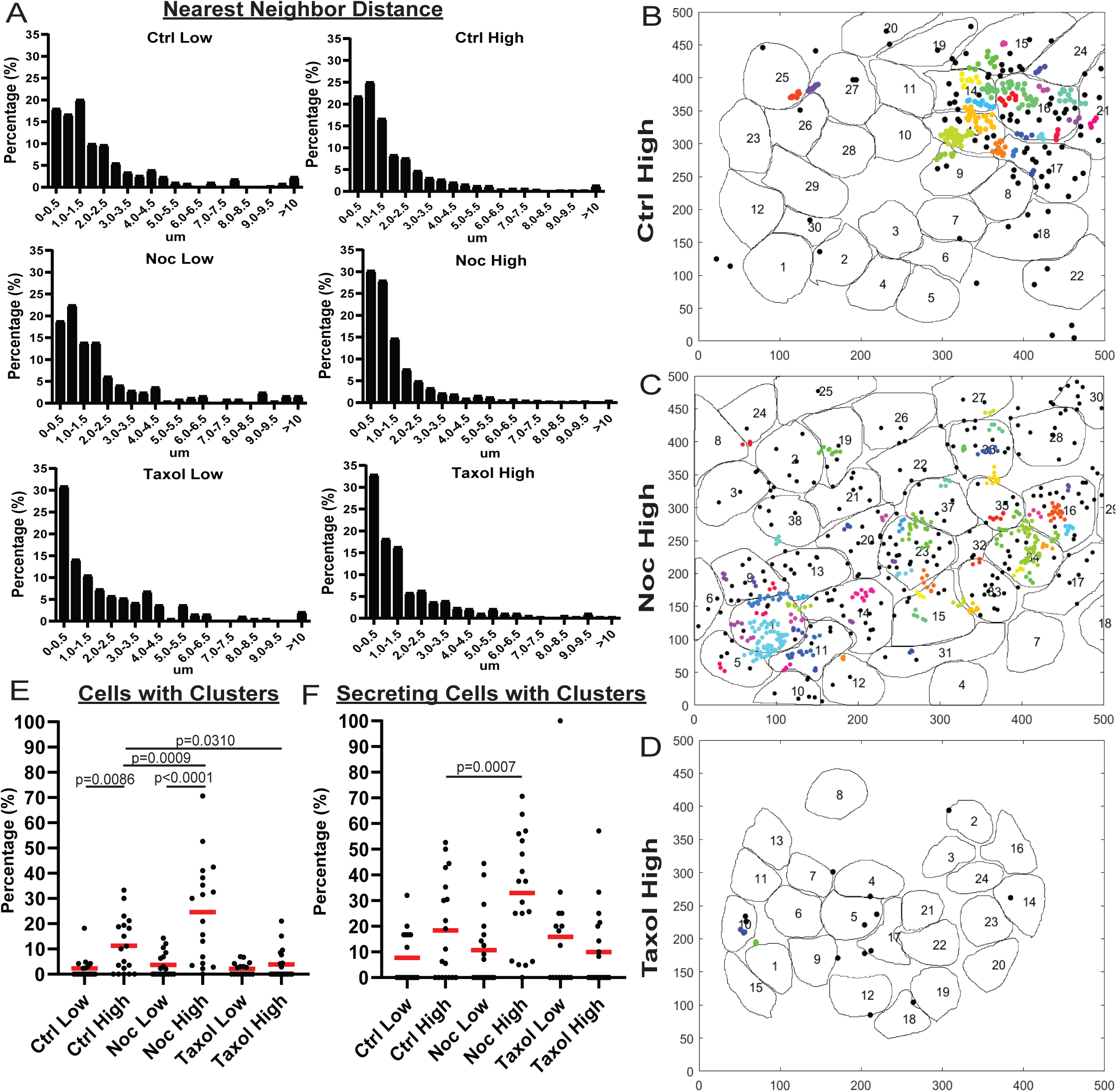
MT stability suppresses formation of insulin secretion hot spots. A) Histogram of nearest neighbor distances obtained by measuring the distance between secretion events in cells with more than one secretion event during the movie. Graphed as percentage of cells within each bin per condition. Bin=0.5 µm. N=191-3255 distances from 16-19 islets. B-D) Representative images of output from Matlab script (see materials and methods) showing cell outlines (black lines) and secretion events (dots). Black dots are non-clustered secretion events, colored dots are clustered secretion events, each different color denotes a different cluster. Clusters were defined as 3+ secretion events occurring within 9 pixels (1.44 µm) by density-based scanning. Islets were pre-treated with DMSO (control, B), nocodazole (C) or taxol (D) and were stimulated with 20 um glucose. E) Graph of the percentage of cells in each field of view with at least one cluster. Red bars, mean. One-way ANOVA and multiple comparison tests, p value as indicated. N= 16-19 islets. F) Graph of the percentage of cells in each field of view that secreted during the movie with at least one secretion event. Red bars, mean. One-way ANOVA and multiple comparison tests, p value as indicated. N= 16-19 islets.

Our analysis (Figure 3E, F) revealed that the number of cells with secretion clusters was small under conditions with low secretion, including all low-glucose treatments and in high glucose after taxol pre-incubation. In contrast, the number of cells with clusters increased from on average 2% to 10% of the cell population by high glucose stimulation in control. Interestingly, nocodazole pre-treatment increased the percentage of cells with clusters in high glucose 2.4 fold (Figure 3E). The nocodazole-induced increase in the number of cells secreting in a non-clustered manner was more modest (1.3 fold). This indicates that in the absence of MTs, a higher proportion of active cells had clustered secretion. Indeed, when only secreting cells were considered, the fraction of cells with clusters increased from on average 18% in control to 33% in nocodazole (Figure 3F). These data suggest that MT depolymerization specifically initiates secretion hot spots to activate dormant β-cells.

### MTs regulate individual cell secretion via both clustered and non-clustered secretion

While our data indicated that the MT-dependent increase in secretion strongly relies on the activation of additional β-cells, we also detected that secretion activity per cell depended on MT presence and stability (Figure 2I, J). Exploring the secretion patterns in individual cells, striking differences between high-glucose stimulated control (Figure 4A, Video 4) and nocodazole-treated (Figure 4B, Video 5) cells are observed. We have found that, in addition to the significant increase in cluster-containing cells, MT depolymerization caused an increase in the number of secretion clusters per cell, even accounting for the increase in secretion (Figure 4C, D). Interestingly, in a vast majority of cells with clustered secretion in control, the number of clusters per cell was restricted to one, while in nocodazole many cells had additional clusters activated (Figure 4E).

**Figure 4.**
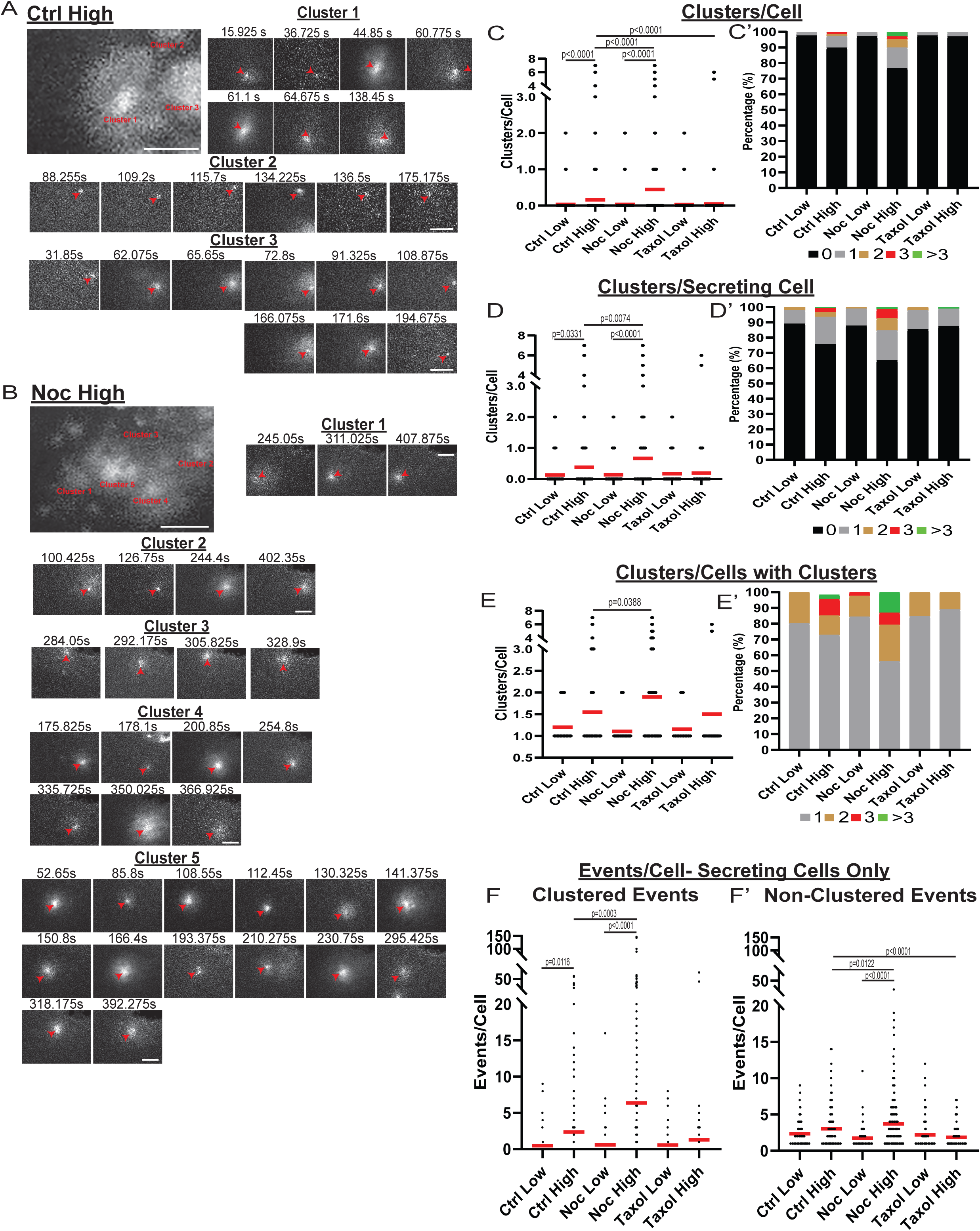
MT-disruption increases the number of hot spots per cell, increasing clustered secretion. A) Representative images of clusters from one cell in a control islet stimulated with 20 mM glucose. Clusters were identified by Matlab script (see materials and methods). First image is time projection through all clusters in the cell 15.925- 194.675 s of the movie, clusters are identified by red text. Time in seconds of each event in the cluster above, red arrowheads identify the secretion event. Scale bar 5 µm. B) Representative images of clusters from one cell in a nocodazole pre-treated islet stimulated with 20 mM glucose. Clusters were identified by Matlab script (see materials and methods). First image is time projection through all clusters in the cell 54.65-407.875 s of the movie, clusters are identified by red text. Time in seconds of each event in each cluster above, red arrowheads identify the secretion event. Scale bar 5 µm. C) Graph of clusters per cell in the field of view, with all cells whether activated during the movie or not. Red bar, mean. Kruskal-Wallis test non-parametric and multiple comparison tests, p value as indicated, N= 495-637 cells from 16-19 islets. C’) Cells from panel C, graphed as a stacked histogram of the percentage of total cells per condition that had each number of clusters. D) Graph of clusters per cell only including cells with at least one secretion event during the duration of the movie. Red bar, mean. Kruskal-Wallis test non-parametric and multiple comparison tests, p value as indicated, N= 88-407 cells from 16-19 islets. D’) Cells from panel E, graphed as a stacked histogram of the percentage of cell with secretion events per condition that had each number of clusters. E) Graph of clusters per cell only cells with at least cluster during the duration of the movie were included. Red bar, mean. Kruskal-Wallis test non-parametric and multiple comparison tests, p value as indicated, N= 13-143 cells from 16-19 islets. E’) Cells from panel D, graphed as a stacked histogram of the percentage of cell with clusters per condition that had each number of clusters. F) Graph of events per cell with at least one secretion event during the movie that were not in a cluster. Red bar, mean. Kruskal-Wallis test non-parametric and multiple comparison tests, N= 88-407 cells from 16-19 islets. F’) Graph of events per cell with at least one secretion event during the movie that were in a cluster. Red bar, mean. Kruskal-Wallis test non-parametric and multiple comparison tests, N= 88-407 cells from 16-19 islets.

In addition, while our data shows that a significant number of exocytic events were specifically concentrated in secretion hot spots, many events were still found randomly scattered across the cell membrane. We tested how the number of non-clustered versus clustered events were changed upon glucose stimulation under conditions of MT depolymerization or stabilization. The number of clustered secretion events per cell increased between low and high glucose, and the loss of MTs resulted in an even higher increase (Figure 4F), reflecting the activation of additional hot spots as indicated above (Figure 4C-E). Interestingly, the non-clustered event number per secreting cell was not significantly increased by high glucose in control, indicating that GSIS is predominantly driven by clustered secretion. At the same time, glucose induced a consistent and significant rise in non-clustered secretion in nocodazole, indicating that without MTs, glucose-stimulated secretion can occur at random, non-hot spot sites (Figure 4F’).

### Individual events within clusters are independent of each other

To better assess how MT stability influences secretion dynamics within hotspots, we analyzed clustered secretion dynamics in comparison to non-clustered secretion. Our analysis detected a slight, statitically insignificant difference in the distribution of cluster sizes (the number of secretion events within a cluster) between conditions (Figure 5A, A’), suggesting that the level of secretion activity within these hot spots is to a large extent MT-independent.

**Figure 5.**
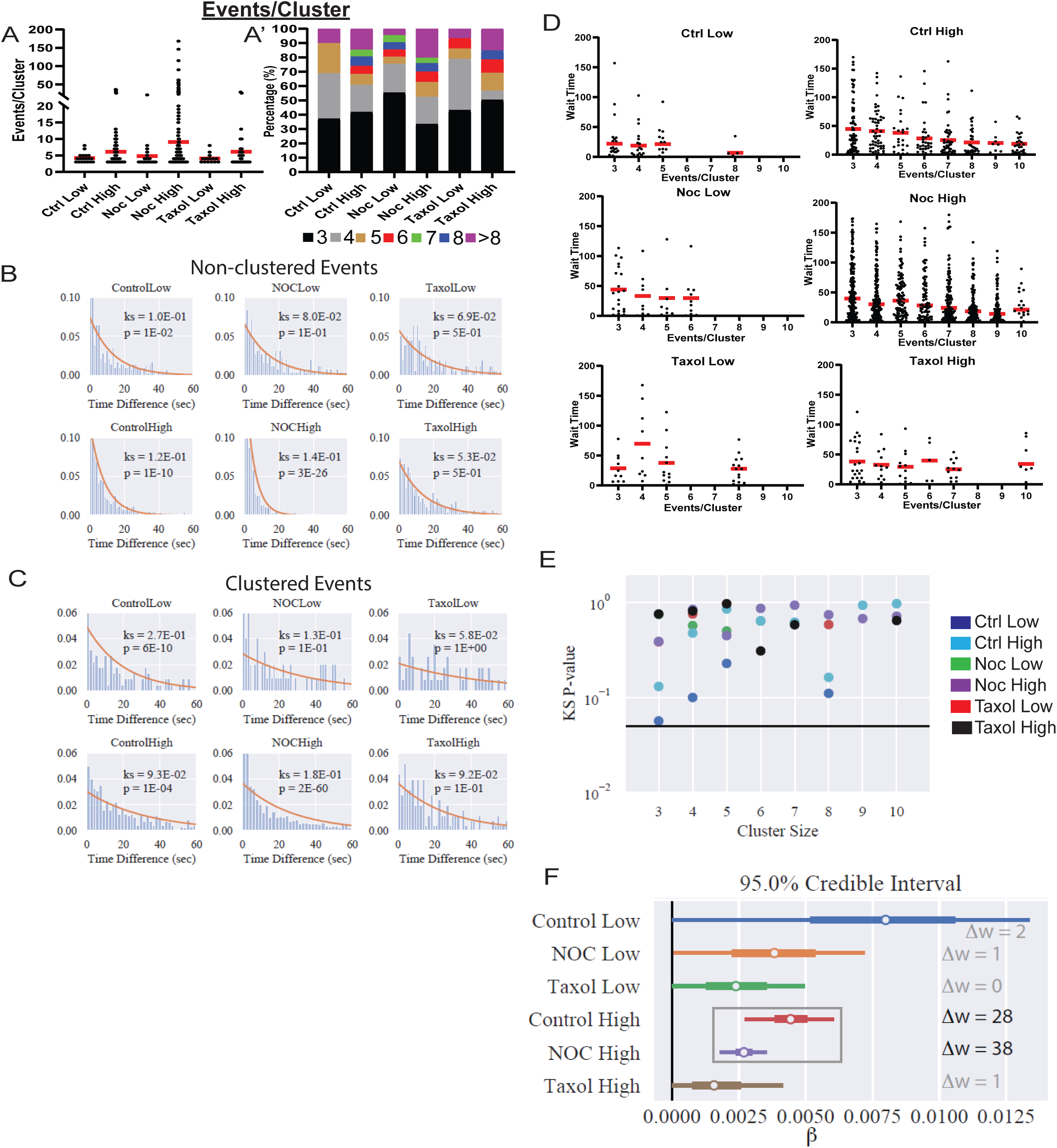
Increased secretion from clusters leads to faster secretion at that site. A) Graph of events per cluster, Red bar, mean. Kruskal-Wallis test non-parametric and multiple comparison tests found no statistical differences between conditions N= 14-290 clusters from 16-19 islets. A’) Clusters from panel F, graphed as a stacked histogram of the percentage of clusters per condition with each number of events. B) Histogram of the time between successively (in time) occurring non-clustered events with the best fit exponential overlaid (KS-statistic is provided for quality of fit). C) Histogram of the time between successively occurring clustered events with the best fit exponential overlaid (KS-statistic is provided for quality of fit). D) Graph of time between successive events distribution for clusters of different sizes (Red bar = mean). Some conditions lack clusters of particular size (e.g. no clusters with 6 secretion events in Ctrl low), as indicated by no data. E) Each distribution in (D) is fit separately to an exponential distribution and quality of fit is assessed with a KS-test (as in panels B, C). The resulting p-value for every test is plotted, with the black line indicating p=0.05, below which the lack of fit is significant. F) Results of fitting a generalized linear model to the data from (D) (see Methods for further details) with the assumption that “secretion rate = α + β * Cluster Size”. Bayesian credible intervals for β are plotted for each condition. This model is also compared to a null model where “secretion rate = α” (i.e. lacking size dependence), with model comparison results reported as the difference of WAIC scores (Δw, positive indicates the full model provides a better accounting of the data).

We next assessed whether clustered and non-clustered secretion events are independent of each other. That is, does one secretion event influence the timing of the next (dependent) or not (independent). To test this, we analyzed the time between events, which if events are independent, should follow an exponential distribution. Non-clustered events are well fit by an exponential distribution (Fig. 5B), suggesting that non-clustered events do not affect each other, as would be expected. A non-parametric Kolmogorov–Smirnov (KS)-test does indicate that, in control and nocodazole treated islets stimulated with high glucose, the distribution does statistically deviate from exponential. However, this is likely due to the clear time dependence of secretion after glucose stimulation in these conditions (Fig. 6A, further discussion below).

**Figure 6.**
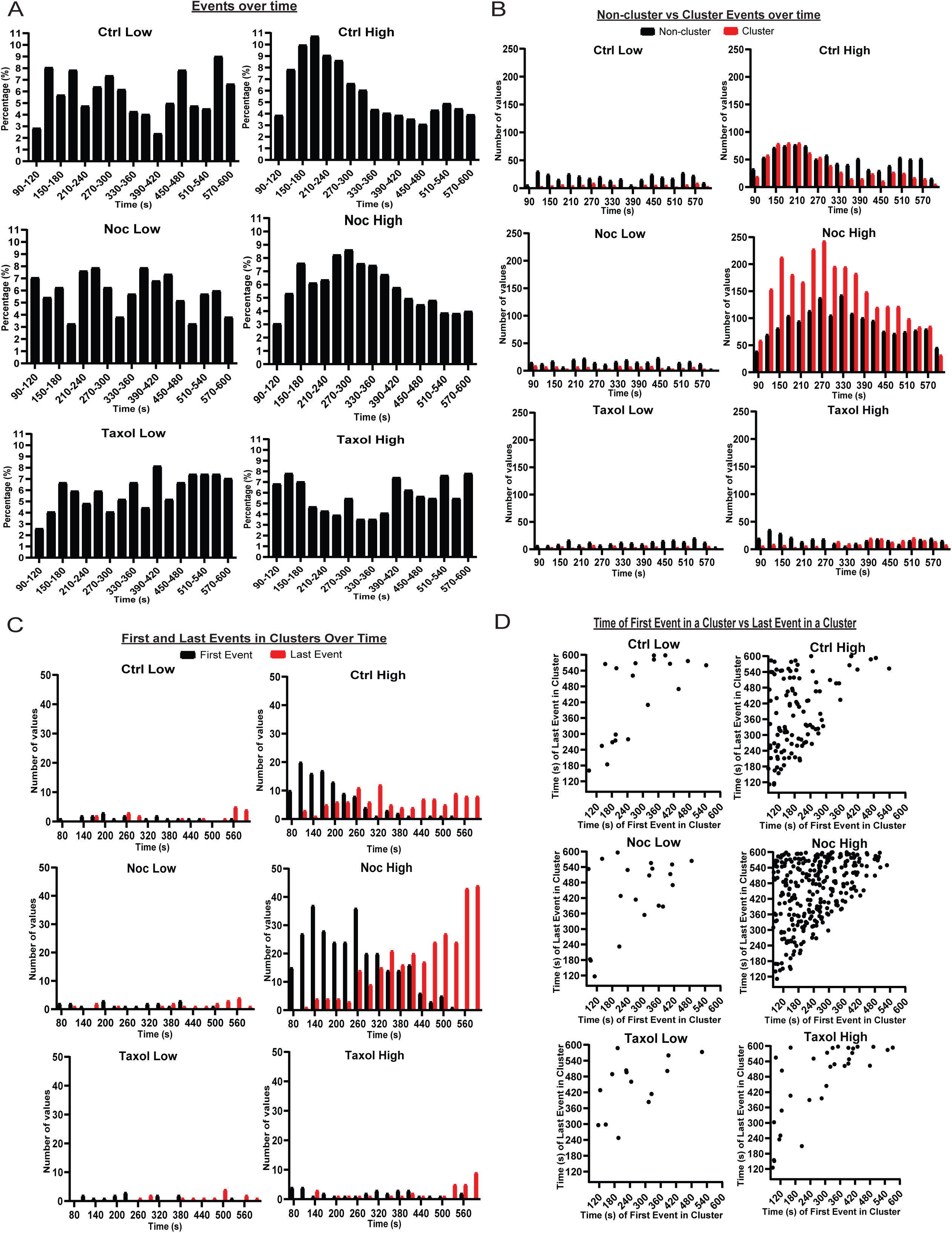
MTs restrict secretion from hot spots to the first phase of secretion, loss of MTs lengthen this phase. A) Histogram of secretion events over time. Graphed as percentage of cells within each bin per condition. Time (seconds) since dye and either high (20 mM) or low (2.8 mM) glucose addition. Bin=30 seconds. N= 16-19 islets. B) Histogram of secretion events over time separated into secretion events not in clusters (black) and in clusters (red). Time (seconds) since dye and either high (20 mM) or low (2.8 mM) glucose addition. Bin=30 seconds. N= 16-19 islets. C) Histogram of the first event in a cluster (black) and last event in a cluster (red) over. Time (seconds) since dye and either high (20 mM) or low (2.8 mM) glucose addition. Bin=30 seconds. N= 16-19 islets. D) Scatterplot of each cluster in each condition with the timing of the first event in a cluster on the x-axis and the timing of the last event in a cluster on the y-axis. Time (seconds) since dye and either high (20 mM) or low (2.8 mM) glucose addition. Bin=30 seconds. N= 16-19 islets.

By comparison, the time-between-events distribution for clustered events clearly deviates from exponential (Fig. 5C). This is particularly obvious under conditions of high glucose stimulation in control and nocodazole, where a high number of clustered events allows for constructing a smooth distribution. Specifically, there is an enrichment of short wait times between events that cannot be captured by the exponential. However, this analysis considers time-between-events within all clusters combined into a single distribution. In this case, the observed deviation may be caused by the presence of clusters with different secretion dynamics within analyzed cluster populations.

To test whether individual clusters exhibit distinct secretion dynamics, we next analyzed event dynamics separately in different cluster sizes. In control and nocodazole treated islets in high glucose (Fig. 5D), the average time between events was lower for larger clusters. Individual analyses of the waiting time distributions for clusters of each size in each of the conditions (Fig. 5E) indicate that those distributions are well fit by exponential distributions. Thus, for each condition, we constructed a generalized linear model (GLiM) with an exponential linking function (since the data are exponentially distributed) where the rate of secretion depends on cluster size according to (*rate =* α + β *\* Size*). This model was fit to the data for each of the six conditions separately using a Bayesian parameter estimation approach.

Our goal is to determine 1) whether (β > *0*) since this would indicate that there is a size dependence of cluster secretion rates and 2) whether the model parameters differ across conditions.

Results of these model fits are presented using 95% Bayesian credible intervals (Kruschke and Liddell, 2018) (Fig. 5F). In brief, a 95% credible interval depicts the range of values that the parameter values will fall into with 95% certainty (note this is technically different than confidence intervals that are often reported in frequentist statistics). Results show that β > *0* in control and nocodazole treated islets in high glucose, indicating that larger clusters do indeed secrete insulin at higher rates. This is confirmed by model comparison where this size-dependent model is compared to a “null” model where the secretion rate is fixed across all sizes (model comparison results are reported using the Watanabe–Akaike information criterion – WAIC, which is a Bayesian companion of the more common AIC). Results are inconclusive in the other four conditions due to the relative scarcity of clustering data leading to more widely distributed credible intervals that overlap with β *= 0*. Comparing model results across conditions shows the credible intervals for both parameters (α not shown) overlap across all six conditions. Thus, there is no discernable difference in the rate of within-cluster secretion, consistent with our results in Fig. 5A.

In conclusion, both clustered and non-clustered events appear to occur independently of each other: one secretion event does not influence the timing of the next. Neither cluster size nor secretion dynamics within clusters differ significantly between any of the examined experimental conditions. However, larger clusters are secreting insulin at faster rates.

### MT stability affects timing of glucose-stimulated insulin secretion

As insulin secretion is known to be tightly regulated in time, we next looked to see if MT stability had any effect on the timing of secretion events. Since the data outlined above show that MT destabilization leads to initiation of extra secretion hot spots, we also analyzed the kinetics of secretion for clustered and non-clustered events separately.

Events in low glucose occurred at random through time with no distinguishable pattern, indicating that basal secretion was random (Figure 6A). Control high glucose showed a more noticeable pattern with the first peak at 2-4 minutes and the second peak at 9-10 minutes after high glucose addition (Figure 6A). Such kinetics correspond well with the known timing of the two waves of the bi-phasic insulin secretion (Curry et al., 1975). Interestingly, we found that clustered secretion at hot spots had a stronger impact on the first phase of secretion, while non-clustered events contributed stronger to the second phase (Figure 6B, Figure 6- figure supplement 1A). In nocodazole-treated islets, high glucose induced a much broader peak of secretion with the maximum at 4-5 minutes, which extended into the timing of the second phase without a defined minimum (Figure 6A). Interestingly, the distribution of non-clustered events maintained a slightly bi-phasic appearance, while clustered secretion appeared as a single wide peak in the time frame of the experiment (Figure 6B, Figure 4- figure supplement 1A). This indicates that timing of clustered secretion was dysregulated.

To test whether the noted distribution is a manifestation of a change in the timing of triggering and/or silencing of secretion hot spots, we analyzed the distribution of the first and last events within each cluster over time (Figure 6C, D). This analysis showed that in control, clusters were initiated mostly in the first 5 minutes after stimulation (first phase), while in the absence of MTs clusters were initiated throughout most of the imaging sequence, suggesting that MTs serve to restrict cluster initiation to the first phase of GSIS (Figure 6C, D).

Furthermore, in control glucose-stimulated islets, over 50% of clustered secretion ceased close to the end of the first phase, while the rest persisted through the second half of the movie. In the absence of MTs, in contrast, a noticeable proportion of clustered secretion lasted through to the end of the recorded sequence, and/or was still active during the last frame (Figure 6C). This indicates that while no difference in the cluster size between control and nocodazole (in any other conditions) was detected, timely cluster inactivation was disturbed in the absence of MTs, and secretion might continue at the same location for a longer period (not recorded due to the fluorescent dye background buildup).

These data are consistent with the model where MTs regulate the timely response of secretion hotspots to the stimulus, including both their initiation and silencing. Lack of such regulation led to “smearing” of the bi-phasic GSIS response, so that the decrease in secretion after the first phase was not evident.

## Discussion

In this study, we have evaluated the role of the MT cytoskeleton in influencing the spatial distribution and heterogeneity of insulin secretion events both at the level of pancreatic β-cell populations and sub-plasma membrane regions within each cell at single cell resolution. Our data indicate that MT stability in the β-cell population is heterogenous and that this heterogeneity is a contributing factor to the previously established heterogeneity of insulin response. This is an exciting finding since functional β-cell heterogeneity has been proposed to allow for an adaptable, efficient response to various changes in physiological conditions and is one of the parameters that dramatically changes in diabetes (Dorrell et al., 2016, Aguayo-Mazzucato et al., 2017). We also show that the loss of MTs causes initiation of additional insulin secretion: 1) activation of hot spots in a higher fraction of cells, 2) increase in the number of hotspots in active cells, and 3) broadens the timing of secretion from the hot spots in the first phase of GSIS. Additionally, MT depolymerization facilitates non-clustered, random secretion activity. Stabilizing MTs prevents GSIS so that all parameters are indistinguishable from basal secretion levels.

There are a number of advantages to the approach utilized in this paper. It is important to measure the spatio-temporal distribution of secretion from individual β-cells in its natural environment in the whole islet with intact cell-to-cell contacts and a proper basement membrane. The major challenge is that this analysis requires TIRF microscopy with β-cells secreting towards the glass coverslip (Loder et al., 2013, Nagamatsu and Ohara-Imaizumi, 2008), however, whole mount islets plated on glass are known to have rapidly compromised responsiveness to glucose. Furthermore, it has become clear that ECM signaling through integrin-dependent calcium channel activity is critical for correct patterning of secretion (Gan et al., 2018, Ohara-Imaizumi et al., 2019b). Our approach of culturing whole mount islets on vascular ECM preserves the functionality of β-cells (Zhu et al., 2015), supporting the idea that the signals downstream of integrins are vital to preserving β-cell identity (Gan et al., 2018). TIRF microscopy in this system allows us to register single secretion events associated with their physiological organizer - the vascular ECM. Thus, this experimental model allows for the evaluation of the secretion patterning arranged by vascular cues, and at the same time, analysis of individual β-cell response as they maintain their connections with each other and other islet cell types. An additional advantage of our approach is that utilization of FluoZin3 dye provides direct information of precise insulin secretion time and location without the need for genetically encoded markers of insulin.

One important conclusion from our study is that MT stability varies in the β-cell population. The mechanistic basis of the differences in MT stability between β-cells is yet unclear. Several possibilities arise from the current literature. For example, since MTs are sensitive to calcium (Hepler, 2016), MT stability might be modified by the calcium influx wave, which is thought to contribute to spatial and temporal differences in β-cell response to stimulation (Benninger et al., 2014). Keeping in mind the evidence that calcium influx is insufficient to provide for secretion heterogeneity (Li et al., 2011), other factors involved in β-cell variability (e.g. cell maturity (Aguayo-Mazzucato et al., 2017, Pipeleers et al., 2017, Gutierrez et al., 2017) gluocokinase expression/activity (Jetton and Magnuson, 1992, Heimberg et al., 1993), islet microenvironment (Trimble et al., 1982, Trimble and Renold, 1981, Brereton et al., 2015), and/or paracrine signaling from other islet cell types (Wojtusciszyn et al., 2008, Pipeleers et al., 1982, van der Meulen et al., 2015, Efendic and Luft, 1975)) could also be involved in regulating MTs. Interestingly, we have shown previously that the β-cell MT network is modified downstream of glucokinase (Zhu et al., 2015, Ho et al, 2020, in press), suggesting that glucokinase variability could lead to heterogeneous MT stability.

One possible important molecular player in this pathway could be the MT stabilizer tau, which we have recently found to be a critical component of glucose- and glucokinase-dependent MT remodeling, leading to efficient insulin release from β-cells (Ho et al, 2020, in press). Other MT-stabilizing proteins found in β-cells, such as MAP2 and doublecortin, may also be involved (Krueger et al., 1997, Jiang et al., 2013). It is clear from our data that regardless of the source of MT heterogeneity, it is functioning to enhance the variability of β-cell secretion activity: pushing MTs toward uniform stability or uniform depolymerization makes β-cell secretion response more uniform in opposite directions. This role of MT heterogeneity in the regulation of variable β-cell response to stimulation is not entirely unexpected. The effect of heterogeneity of MT stability in cell populations has been described in other cell types and is implicated in the variability of cell function. For example, motile cells that have directionally stable MT arrays have an increased ability for migration in wound healing assays (Sugioka and Sawa, 2012). Heterogeneity of MT dynamics in neurons has been found to underlie the distinction between the axon and dendrites (Conde and Caceres, 2009), neurite capacity for cargo movement (Franker and Hoogenraad, 2013), and modulation of local signaling and rearrangements in neuronal connectivity (Hoogenraad and Bradke, 2009). Now, our data add β-cells to the list of cell types where the differential MT stability plays an important role in cell physiology.

How do differential MT dynamics promote differential secretion? The first possibility is the direct MT regulation of insulin secretion, when differently organized MT networks differentially transport insulin granules within individual cells. This aligns well with the model supported by our previous findings (Zhu et al., 2015, Bracey et al., 2020, Ho et al, 2020, in press) where stable MTs act as tracks for the withdrawal of insulin granules from the plasma membrane, restricting secretion. Lack of this regulated withdrawal should allow for the increased secretion at random locations. Indeed, this is what we observed: the increase of glucose-stimulated non-clustered events when MTs are absent (nocodazole treatment). MT-dependent withdrawal could also contribute to the restriction of clustered secretion if secretion-competent granules are specifically accumulated near the hot spots by a potential additional mechanism.

The other possibility is that MTs act indirectly by modulating one of the prominent mechanisms involved in β-cell heterogeneity regulation. Interestingly, such master regulators of heterogeneous β-cell response such as gap junctions and ion channels have been shown to be modulated by MTs in other cell types (Shaw et al., 2007). A scenario where MTs are both regulated by calcium and tune calcium influx in a feedback loop has been discussed previously, for example in immune cells (Joseph et al., 2014). It is interesting in this regard that we found that MTs regulate heterogeneity of β-cells responsiveness to glucose specifically via activation of secretion clusters, or hot spots. Existing evidence proposes that loci of repeated clustered secretion could depend on membrane localization of VDCC (Bokvist et al., 1995), secretion machinery (SNARES, etc) (Ohara-Imaizumi et al., 2007, Ohara-Imaizumi et al., 2004), K+ channels (Fu et al., 2019), and ELKS (Ohara-Imaizumi et al., 2005, Ohara-Imaizumi et al., 2019a). ELKS is a particularly interesting molecular player, since accumulating evidence implicates it in local activation of L-type calcium channels and site-specific calcium influx (Ohara-Imaizumi et al., 2019b). ELKS is thought to be responsible for directional insulin secretion toward vasculature, because it accumulates on the β-cell membrane at the sites of vascular contact (Ohara-Imaizumi et al., 2005) downstream of integrin activation by vascular ECM proteins (Gan et al., 2018, Low et al., 2014). At the same time, extensive literature connects MTs with regulation of integrin based ECM adhesions (focal adhesions) in a variety of cell types: MT depolymerization promotes assembly, and MT regrowth facilitates adhesion turnover (Kaverina et al., 1999, Stehbens and Wittmann, 2012, Ezratty et al., 2005). In Hela cells, assembly of adhesion-associated patches of ELKS and its binding partner LL5B is also facilitated by MT depolymerization (Lansbergen et al., 2006). In β-cells, a similar mechanism driven by MT destabilization may lead to the formation of vascular-ECM dependent patches on the membrane where clustering of secretion events can occur. Thus, clustered secretion could be regulated through this molecular link by a combination of vascular ECM and MTs.

Another possibility is that a reduction in MT stability regulates local calcium influx via STIM-1, a calcium transporter that was shown to be positioned and regulated by dynamic MTs via its binding to MT plus end binding protein EB1 (Tomas-Martin et al., 2015, Usui et al., 2019). An additional scenario is that dynamic MTs facilitate secretion at hotspots by coupling insulin exocytosis with compensatory endocytic events as was shown for ELKS/LL5beta patches in cultured Ins-1 cells (Yuan et al., 2015a).

Regardless of the mechanism, the important conclusion from our data is that it is the activation of hot spots in additional cells that makes the β-cell population response more uniform in the absence of MTs. Since calcium is essential but likely insufficient for β-cell heterogeneity (Benninger and Hodson, 2018), it is possible that suppression of hot spot assembly by stable MTs provides a required layer of control restricting secretion in a subpopulation of cells in the presence of calcium influx.

Another important take-home message from our analyses is that MTs regulate the timing of the first phase of GSIS via control of clustered secretion. We show that initiation of secretion hot spots that is normally rapidly triggered by the metabolic signal is dysregulated when MTs are lost. In the absence of MT regulation, new secretion sites continue to appear even at the time points when the first phase of secretion should normally be fading. This suggests a potential role for MTs in control of the end of the first phase of GSIS, which could be a significant part of prevention of over secretion under physiological conditions. At the same time, MT-dependent regulation of random secretion applies to both stages of the bi-phasic GSIS. An extended first phase of secretion and enhanced secretion levels in both phases are consistent with our previous findings (Zhu et al., 2015).

At the same time, we do not find a significant role of MTs in the secretion dynamics within individual clusters. Our data clearly indicate that MTs do not contribute into the frequency of secretion events within clusters, which is at odds with a previous finding in KCl-stimulated Ins-1 cells plated on glass, where a decreased frequency of events within a cluster was observed in the absence of MTs (Yuan et al., 2015b). It is unclear at this point whether this discrepancy is due to the dramatic differences in the experimental models used in the two studies, but it is not surprising that the fine regulation detected here was not observed in the absence of islet cell interactions and vascular ECM signals.

## Materials and Methods

### Mouse utilization

Ins-Apl mice with Histone 2B-mApple knocked into the Ins2 locus (Stancill et al., 2019) were used for all experiments. Males and females between 2-6 months were used. Data was separated by sex to determine statistical differences before being combined. Mice utilization was supervised by the Vanderbilt Institutional Animal Care and Use Committee (IACUC).

### Cell lines and maintenance

RPE1-hTert (ATCC) cells were maintained in DMEM/F12 with 10% FBS and antibiotic at 37 °C with 5% CO2 and were periodically tested for mycoplasma.

### Islet picking and dissemination

Mouse pancreatic islets were hand-picked following *in situ* collagenase perfusion and digestion. Islets were allowed to recover for at least 1 hour in RPMI 1640 Media (Life Technologies, Frederick, MD) supplemented with 10% fetal bovine serum (FBS), 100 U/mL penicillin, and 0.1cmg/mL streptomycin in 5% CO_2_ at 37°C.

For experiments utilizing intact islets, 5-8 islets per 10 mm glass bottomed dish (Mattek) were placed in the center of the glass in 100 ul RPMI 1640 and allowed to settle for 1.5-2 hours before being transferred to 5% CO_2_ at 37°C. The following day, 900 ul of RPMI 1640 was added. Islet media was changed every 2-3 days for 5-9 days when the experiment was performed.

For experiments utilizing disseminated islets, 30-50 islets per coverslip or dish were used. Islets recovered for 4-24 hours following picking. To disseminate, all islets were collected into a 15 ml tube on ice and allowed to settle for 5 minutes. Most of the media was removed and islets were resuspended in 900 ul room temperature versine and mixed by pipetting up and down several times, and 100 ul warm 0.05% trypsin-EDTA was added. The mixture was pipetted up and down 25-30 times with a p1000 tip to allow for dissemination but keep clumps of islet cells present. Disseminated islets were then centrifuged at 300 x G for 2 minutes at room temperature. The versine/trypsin solution was removed and cells were resuspended in 100 µl RPMI 1640 media per 30-50 islets. The cell mixture was added to coverslips or dishes.

All coverslips and dishes were plasma cleaned and coated in placenta-dereived human ECM (Corning, cat#: 354237) which is comprised of laminin, collagen IV and heparan sulfate proteoglycan which resembles vasculature ECM for 30 minutes at 37°C. For disseminated islet cells used in Fluozin-3 assays, dishes were scratched before plasma cleaning with a diamond pen in a pattern to aid in finding the same cells after live cell imaging and immunostaining

### Fluozin-3 assay

Fluozin-3 assay was preformed 5-9 days after picking to allow for flattening of the islets and the day following dissemination for disseminated islets. 16-20 hours prior, RPE1-hTert cells stably expressing Centrin-GFP were plated with islets at <5% confluency. The green signal in these cells is diffuse outside of the centrosome signal. These cells allow for more accurate determination of focus and TIRF angle for the assay as the green Fluozin-3 signal only appears once it is added to the dish.

On the day of the assay, islets were incubated at 37°C in low glucose (2.8 mM glucose) KRB (110 mM NaCl, 5 mM KCl, 1 mM MgSO4, 1 mM KH2PO4, 1 mM HEPES, 2.5 mM, CaCl2 and 1 mg/mL BSA) for 1.5-2 hours with a change of buffer after 1 hour. For nocodazole (Sigma-Aldrich Cat#: M1404) treatment, stock solution was added to a final 5 µM concentration to treat islets for at least 4 hours before imaging. For taxol treatment (Sigma-Aldrich Cat#: PHL89806), islets were treated for two hours before imaging with 5 µM taxol. Immediately before imaging the buffer was replaced with 100 ul fresh buffer with the same treatments as before.

Dishes were then placed on the TIRF microscope and allowed to equilibrate for 10 minutes. An islet was identified by eye. A nearby Centrin-GFP RPE cell was used to focus the microscope to the bottom of the dish and set the TIRF angle of the green laser. A 10 µm stack of 0.2 µm slices was recorded of the islet before addition of the dye using both transmitted light and the 568nm laser. Stacks were started below the islet to ensure the bottom of the cells were imaged. 50 µl of KRB buffer with the Fluozin-3 dye to final concentration of 20 µM was added. For high glucose treatment glucose to a final concentration of 20 µM was added, after which the imaging sequence was started (60ms exposure, no delay for 10 minutes). Focus and TIRF angle were refined after dye addition.

### Processing of Fluozin-3 movies

The first 1,600 frames (approximately 1.8 minutes) from each movie were removed as the focus and TIRF angle were being adjusted during this time and the dye needs time to penetrate. Each image was then subtracted from the previous image, briefly the first and last frame from two different copies of the file were removed and subtracted using the Image Calculator tool in ImageJ with the 32-bit (float) result box checked. Every 5 images from this subtracted image were grouped as a max projection through time using the Grouped Z-project function in ImageJ.

### Analysis of Fluozin-3 movies

#### Identifying secretion events

Individual secretion events or “flashes” were identified using ImageJ software (Schindelin et al., 2012). Fluozin-3 movies were blinded then analyzed manually. A macro was written that identifies the centroid of local maximal signals after the users uses the point function identify a bright signal. Each event was identified as a flash of signal that was present in one frame only, with a noise level of 5000 and a search range of 6 pixels. Centroid coordinates were exported and used for analysis. On occasion bright spots in the center of the event caused the formation of donuts preventing accurate identification of the centroid. These rare events were noted during analysis and the centroid was identified manually and added to the data.

#### Identifying cell borders

Transmitted light and 568nm stacks were recorded before dye addition to identify individual β-cell borders. All cells with a red nucleus (Ins-Apl signal) were outlined by hand in ImageJ. If the nuclei could not be seen (above the image stack range, signal diminished because of light dispersal or out of the frame) or was Ins-Apl negative, the cell was assumed to be a non-β-cell and discarded from analysis. Each Ins-Apl positive cell outline was saved as an individual ROI in ImageJ and coordinates were exported.

#### Identifying secretion events/cell and clusters

A Matlab script (see supplemental annotated scripts) was used to compare the location of each secretion event (identified above) in relation to each cell (borders identified by ROI), outputting the number of secretion events in each cell. Some secretion events fell outside the boundaries of all β-cells identified and were discarded from analysis. Within the same annotated Matlab script clustering analysis was performed. Density based scanning was used with a neighborhood search radius of 1.5 µm and a minimum number of neighbors of 3. The script outputs (1) the movie frame (time), centroid, cell and cluster of each identified secretion event, (2) the cell number, secretion events and clusters within each identified cell, and (3) the size of each cluster.

### Immunofluorescence

#### Intact attached and disseminate islets following Fluozin-3 assay

Dishes were removed from the microscope stage and washed 5 times in PBS to remove the Fluozin-3 dye. Islets were fixed in 4% paraformaldehyde in PBS with 0.1% Triton-X 100 and 0.25% Glutaraldehyde for one hour (intact islets) or ten minutes (dissemintated islets) at room temperature. Following fixation, islets were washed 5 times in PBS with 0.1% Triton-X 100 at room temperature.

#### Disseminated islets for tubulin and Glu-tubulin intensity measurement

Dishes or coverslips were pre-treated with low glucose (2.8 mM) KRB at 37°C for 1.5-2 hours with a change in buffer after an hour. High glucose dishes or coverslips were then treated with high glucose (20 mM) KRB for 10 minutes. For ice treatment, the dishes or coverslips were placed on ice for 30 minutes before fixation. For cytosolic pre-extraction (dilution), cells were place in 0.1% Triton in PEM buffer (0.1 M PIPES (pH 6.95), 2 mM EGTA, 1 mM MgSO_4_) for one minute, then Triton was washed out and islets were kept at 37°C in PEM buffer for 20 minutes before fixation. Cells were then fixed in 4% paraformaldehyde in PBS with 0.1% Triton-X 100 and 0.25% Glutaraldehyde. Following fixation, islets were washed 5 times in PBS with 0.1% Triton-X 100.

Primary and secondary antibodies were incubated for 24 hours (disseminated islets) or 48 hours (intact islets, antibody change after 24 hours). Samples were washed three times in PBS + 0.1% Triton-X 100 after primary and secondary antibody incubations. Following the final wash post-fixing in 4% paraformaldehyde was performed (30 minutes for intact islets and 10 minutes for disseminated islets) and one more round of washing was performed before mounting coverslips.

Primary antibodies used are mouse anti-α -tubulin, DM1A clone (1:500, dilution, Sigma-Aldrich, Cat#: T9026), rabbit anti-detyrosinated tubulin (1:500, dilution, Millipore, cat#: AB3201). Alexa488- and Alexa647-conjugated highly cross-absorbed secondary antibodies were obtained from Invitrogen. Coverslips were mounted in Vectashield Mounting Medium (Vector Labs Cat#: H-1000).

### Microscopes

Fixed samples were imaged on a laser scanning confocal microscope Nikon A1r based on a TiE Motorized Inverted Microscope using a 100X lens, NA 1.49, run by NIS Elements C software. Cells were imaged in 2 µm slices through the whole cell for disseminated islets. Intact islets were imaged through 20 µm. Images in Figure 1 are single slices from the bottom of the cells. Image in Figure 1- figure supplement 1 are maximum intensity projections across three slices from the bottom of the islet to better show the MT cytoskeleton.

Fluozin-3 assays for secretion analysis were imaged on a Nikon TE2000E microscope equipped with a Nikon TIRF2 System for TE2000 using a TIRF 100× 1.49 NA oil-immersion lens and an Andor iXon EMCCD camera run by NIS Elements C software.

### Analysis of average intensity of Tubulin and Glu-tubulin

β-cells were identified using red nuclei and outlined with each β-cell being assigned an ROI. The bottom of the cells where used for analysis as cell boarders were clearly visible and secretion in this location was measured in Fluozin-3 assays. Using ImageJ, the mean intensity of each cell in one slice at the bottom of the stack was measured. Both the Glu-tubulin and Tubulin channels were measured. Bright primary cilia signal was removed through thresholding of bright signals, assigning an ROI and deleting the signal from both channels. A small box was drawn outside of the cell and measured in both channels for each image as the background. Background was then subtracted from the mean intensity within the cell.

### Statistical modeling methods

Given the distributional nature of this secretion event data, we use statistical modeling to both assess the properties of the process giving rise to the observed data and determine how those properties change under different conditions. A natural hypothesis to test is that secretion events occur independently of each other. If events are indeed independent, then the time between successive events within a cell or cluster should be exponentially distributed. To test this, we fit each data set using an exponential distribution using python’s SciPy package. For non-cluster data, all events occurring within a cell were grouped in ascending order of occurrence time to produce a time-between-event distribution that was fit to an exponential distribution. For analysis of all cluster data within each condition (e.g. high glucose Taxol treated), events occurring within each cluster were grouped to compute the time between successive events within the same cluster. These wait times for all clusters were then grouped together to form a time-between-event distribution that was fit. This analysis however grouped clusters of different sizes together. We thus separately analyzed clusters of different sizes (measured as secretion events per cluster). For this, we collected the within cluster time between events for clusters of each size separately. We then grouped all computed waiting times for clusters of a given size (e.g. 4 secretion events) and fit the resulting distribution for each size separately to an exponential.

To assess whether clusters of different sizes secrete insulin at different rates, we constructed a generalized linear model (GLiM) where the secretion rate is linearly dependent on cluster size. Since the secretion time data for each cluster size is well approximated by an exponential distribution (as verified by the previously described analysis), we use an exponential linking function. Note that due to the non-normal nature of this data and the relatively small sample sizes, this GLiM approach is more appropriate than a more common ANOVA or similar approach. This GLiM was fit to the data for each condition separately using Bayesian parameter estimation with the python PyMC package. Both the parameters (□, □) were assigned half-normal priors with a standard deviation of 0.1, which are weakly informative priors. Four MCMC chains were used with 4,000 samples each using the built in NUTS sampler.

### Experimental Design

Glu-tubulin and tubulin intensity experiments were replicated three times. At least 20 images per experiment were obtained, random fields containing β-cells (identified by red nuclei) were imaged.

Fluozin-3 assay experiments were repeated on at least seven different days. If at least one secretion event was identified, the movie was analyzed. Any movies without a secretion event were excluded as the technical difficulty of the assay caused an inability to identify a reason for lack of secretion. Analysis of secretion events was performed blind.

After analysis no data was excluded from any experiment.

### Image Processing

In order to make small structures visible, adjustments were made to brightness, contrast and gamma settings of all fluorescent images presented here.

### Statistics

For data sets where the distribution was appropriate (as determined by the D’Agostino & Pearson omnibus normality test), statistics were calculated by one-way ANOVA with Tukey’s multiple comparisons test. When the data was not normal, a non-parametric test was used, Kruskal-Wallis test non-parametric and multiple comparison tests. GraphPad Prism was used for statistical analyses and graphical representations. Significance was defined at p ≤ 0.05.

## Abbreviations

(GSIS): Glucose-stimulated insulin secretion
(MT): Microtubule
(Noc): Nocodazole
(GLiM): Generalized Linear Model
(ECM): Extracellular matrix
(TIRF): Total Internal Reflection Fluorescence
(KS): Kolmogorov–Smirnov

## Acknowledgements

This work was supported by National Institutes of Health (NIH) grants T32 DK07061 and 1F32DK117529 (to K.P.T.), R35-GM127098 (to I.K.), R01-DK65949 (to G.G.), DMS1562078 (to W.R.H.), and R01-DK106228 (to I.K. and G.G.). and R35-GM119552 (to M.Z.). M.Z. acknowledges support from the Searle Scholars Program. We thank Hamida Ahmed, Dr. Alexey Khodjakov and Dr. Anastasia Kaverina for technical help. The authors also thank Dr. Anneke Sanders for critical reading of the manuscript.

## Author Contributions

K.P.T. performed all experiments and data analysis and wrote the manuscript. H.K., G.A. T.G.F. and M.Z. provided scripts for data analysis. Z.X. provided preliminary data and performed troubleshooting of experimental methods. A.B.O and M.A.M provided materials. G.G. Provided conceptural insight and wrote the manuscript. W.R.H. provided mathematical modeling, provided conceptual insight and wrote the manuscript. I.K. supervised the study, provided conceptual insight and wrote the manuscript.

## Declation of Interests

The authors declare no competing interests.

## Figure Legends

**Figure 1- figure supplement 1. MT stability is regulated by glucose stimulation.**

A) Scatterplot of tubulin average intensity for each cell. Mean, red bar. n=101 cells per condition.

B) Histogram of tubulin average intensity in low (black) and high (red) glucose. Bin=100. n=101 cells per condition, Students’ t-test found no statistical significance.

C) Histogram of tubulin average intensity normalized to the mean of each low (black) and high (glucose). Bin=0.2. Coefficient of variation= standard deviation/mean. n=101 cells per condition.

D-E) Glu-tubulin staining in disseminated islets after 30 min on ice (corresponds to Figure 1M-O). β-cells outlined in dashed yellow lines. Single slice from the bottom of the cell. Scale bar 10 µm.

F) Scatterplot of Glu-tubulin average intensity for each cell after 30 minutes in high glucose. Mean, red bar. Students’ t-test, p<0.0001. n=100-101 cells per condition

G) Histogram of Glu-tubulin average intensity in low (black) and high (red) glucose after 30 min on ice. Bin=100. n=100-101 cells per condition

H) Histogram of Glu-tubulin average intensity normalized to the mean of each low (black) and high (glucose) after 30 min on ice. Bin=0.2. Coefficient of variation= standard deviation/mean. n=100-101 cells per condition.

I) Scatterplot of Glu-tubulin to tubulin ratio of average intensity for each cell after 30 min on ice. Mean, red bar. Students’ t-test, p=0.135. n=100-101 cells per condition.

J) Histogram of Glu-tubulin to tubulin ratio of average intensity in low (black) and high (red) glucose after 30 min on ice. Bin=1.0. n=100-101 cells per condition

K) Histogram of Glu-tubulin to tubulin ratio of average intensity normalized to the mean of each low (black) and high (glucose) after 30 min on ice. Bin=0.2. Coefficient of variation= standard deviation/mean. n=100-101 cells per condition

L-N) Disseminated islets placed extracted for one minute and placed in buffer for 20 minutes. Stained for Glu-tubulin (L) and tubulin (M). β-cells were identified by red nuclear Ins-Apl expression (N, yellow), merged with Glu-tubulin (cyan) and tubulin (magenta). Red arrows pointing to differences between cells. Single slice from the bottom of the cells. Scale bar 10 µm.

**Figure 2- figure supplement 1. Assay protocol and basal glucose conditions are not affected by MT stability. Corresponds to Figure 2.**

A) Overview of Fluozin-3 assays. For more information see materials and methods.

B-D) Time projections of islets from Videos 1-3 (low glucose) inverted. Fluozin-3 flashes are represented as black areas. Cell borders overlaid in red. Islets were pre-incubated in DMSO (control, B), nocodazole (C) or taxol (D) and incubated in 2.8 mM glucose. Scale bars 100 µm.

E) Cell outlines (white line) and secretion events (red circles) from a disseminated islet after 10 minutes in 20 mM glucose and Fluozin-3 dye. Scale bar 10 µm.

F-G) Disseminated islet from E post-fixed following TIRF imaging for Glu-tubulin (F) and tubulin (G). Cells (yellow dashed lines) correspond to cells in E with the same number. Singe slice from the bottom of the cells. Scale bar 10 µm.

H-J) Representative images of tubulin following Fluozin-3 imaging as shown in A, islets were pre-incubated in DMSO (control, H), nocodazole (I) or taxol (J) and stimulated with 20 mM glucose. 3-image max projection of the bottom of the islet. Scale bars, 10 µm.

**Figure 3- figure supplement 1. 3+ event clusters are not spurious and a very rare in basal glucose conditions. Corresponds to Figure 3.**

A) Computationally simulated random secretion events in *in silico* cells using the mean area of the cells analyzed.

B-D) Representative images of output from Matlab script (see materials and methods) showing cell outlines (black lines) and secretion events (dots). Black dots are non-clustered secretion events, colored dots are clustered secretion events, each different color denotes a different cluster. Clusters were defined as 3+ secretion events occurring with 9 pixels (1.44 µm) by density-based scanning. Islets were pre-treated with DMSO (control, B), nocodazole (C) or taxol (D) and were incubated in 2.8 um glucose.

**Figure 6- figure supplement 1. Clustered secretion is mostly restricted to the first phase of secretion in control islets. Corresponds to Figure 5.**

A) Histogram of secretion events over time separated into secretion events not in clusters (black) and in clusters (red). Graphed as percentage of cells within each bin per condition. Time (seconds) since dye and either high (20 mM) or low (2.8 mM) glucose. Bin=30 seconds. N= 16-19 islets.

## Movie Legends

**Video 1. Control islet insulin secretion in low and high glucose. Related to Figure 2.**

DMSO treated islets in low glucose (left) and high glucose (right). Five frame projection through time, each slice is 325 ms over about 8.5 minutes. Fluozin-3 dye creates flashes upon zinc binding, representing a single insulin secretion event.

**Video 2. Nocodazole islet insulin secretion in low and high glucose. Related to Figure 2.**

Nocodazole treated islets in low glucose (left) and high glucose (right). Five frame projection through time, each slice is 325 ms over about 8.5 minutes. Fluozin-3 dye creates flashes upon zinc binding, representing a single insulin secretion event.

**Video 3. Taxol islet insulin secretion in low and high glucose. Related to Figure 2.**

Taxol treated islets in low glucose (left) and high glucose (right). Five frame projection through time, each slice is 325 ms over about 8.5 minutes. Fluozin-3 dye creates flashes upon zinc binding, representing a single insulin secretion event.

**Video 4. Clusters in a single cell in control islet in high glucose. Related to Figure 4.**

A single cell from Video 1 high glucose. Projection through time of 5 frames, each slice is 325 ms. Movie is 15.925-194.675 s of the movie.

**Video 5. Clusters in a single cell in nocodazole treated islet in high glucose. Related to Figure 4.**

A single cell from Video 2 high glucose. Projection through time of 5 frames, each slice is 325 ms. Movie is 54.65-407.875 s of the movie.

