## Supplementary figures and images for "Microtubules regulate pancreatic beta cell heterogeneity via spatiotemporal control of insulin secretion hot spots"

### Figure 1- figure supplement 1

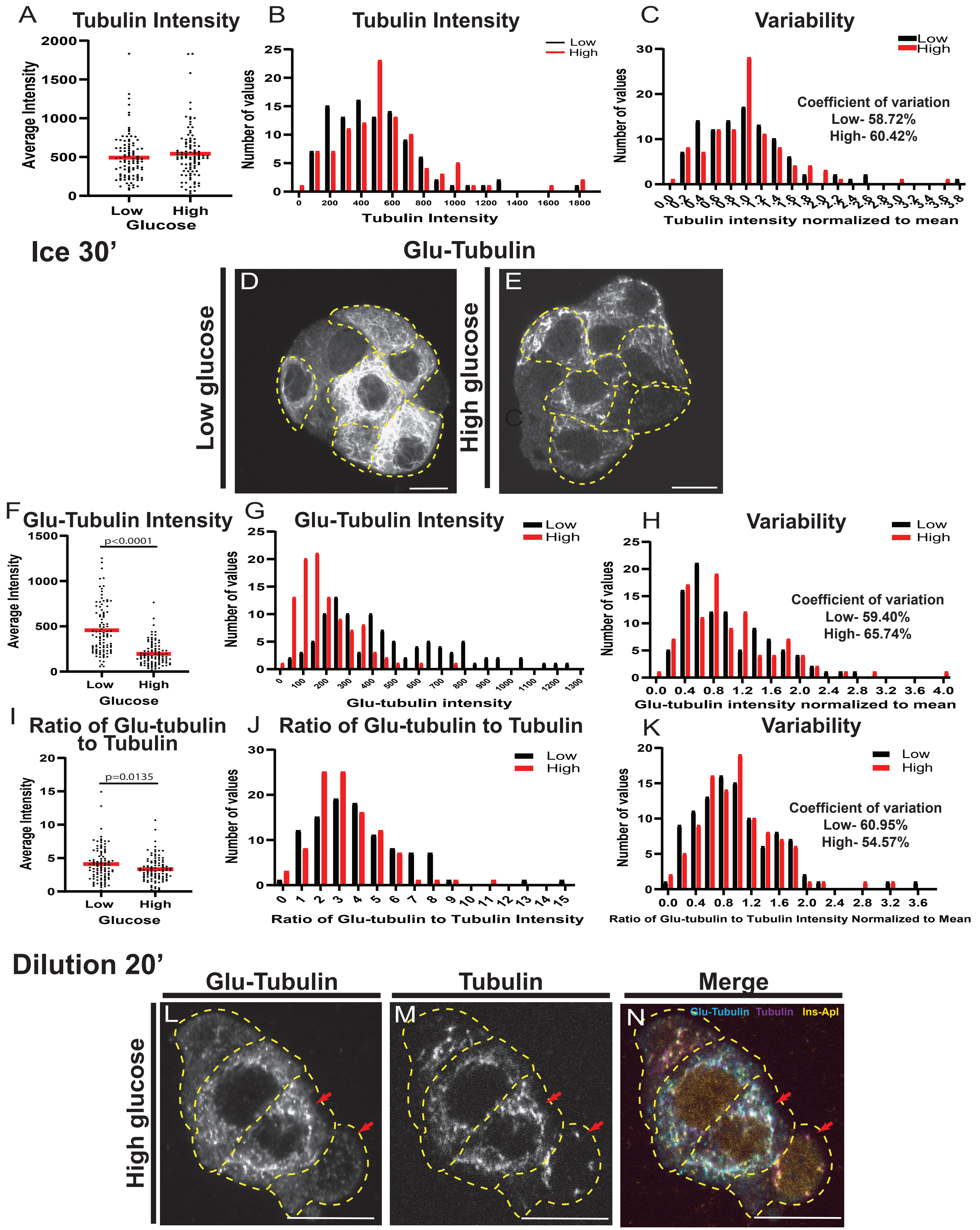

### Figure 2- figure supplement 1

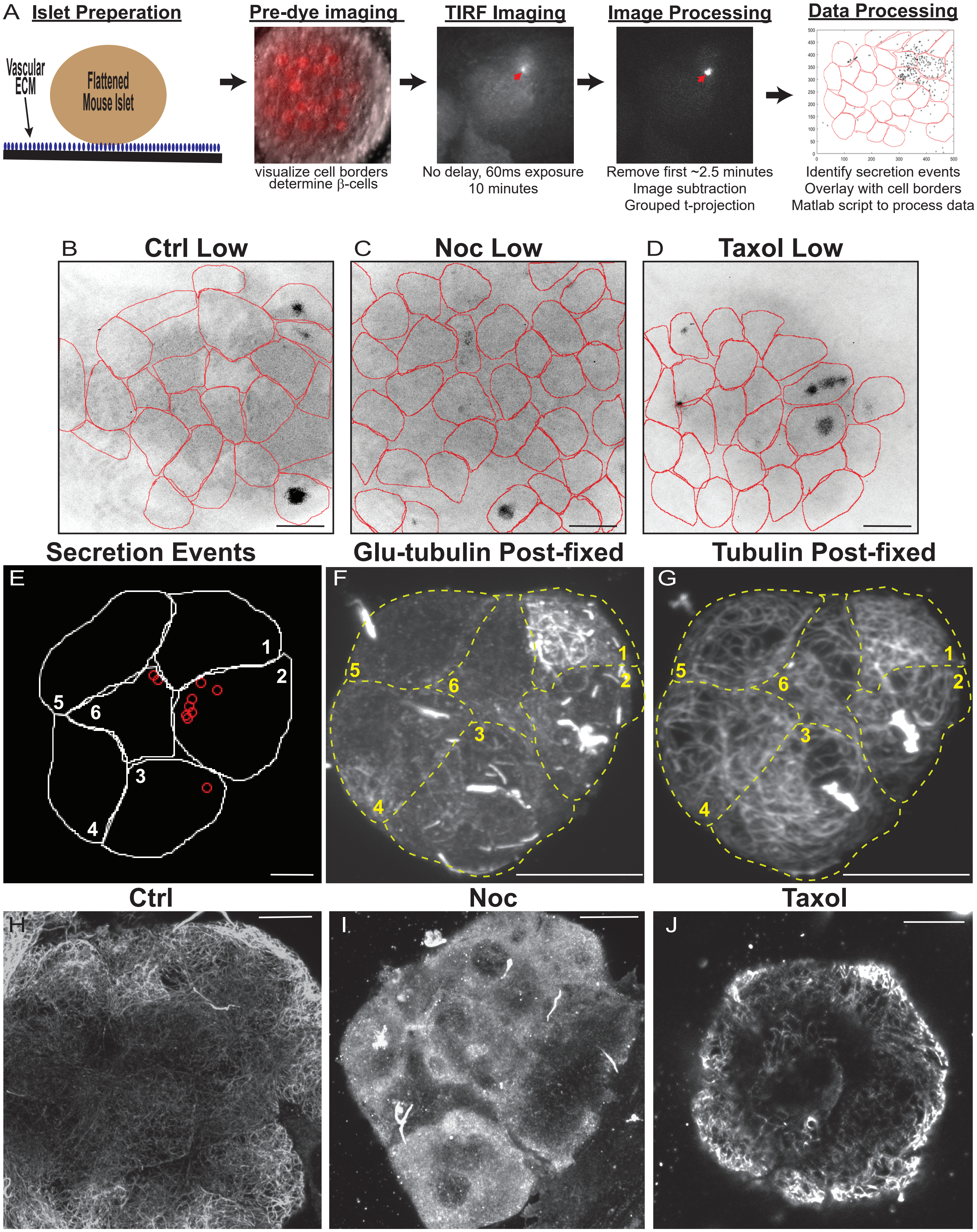

### Figure 3- figure supplement 1

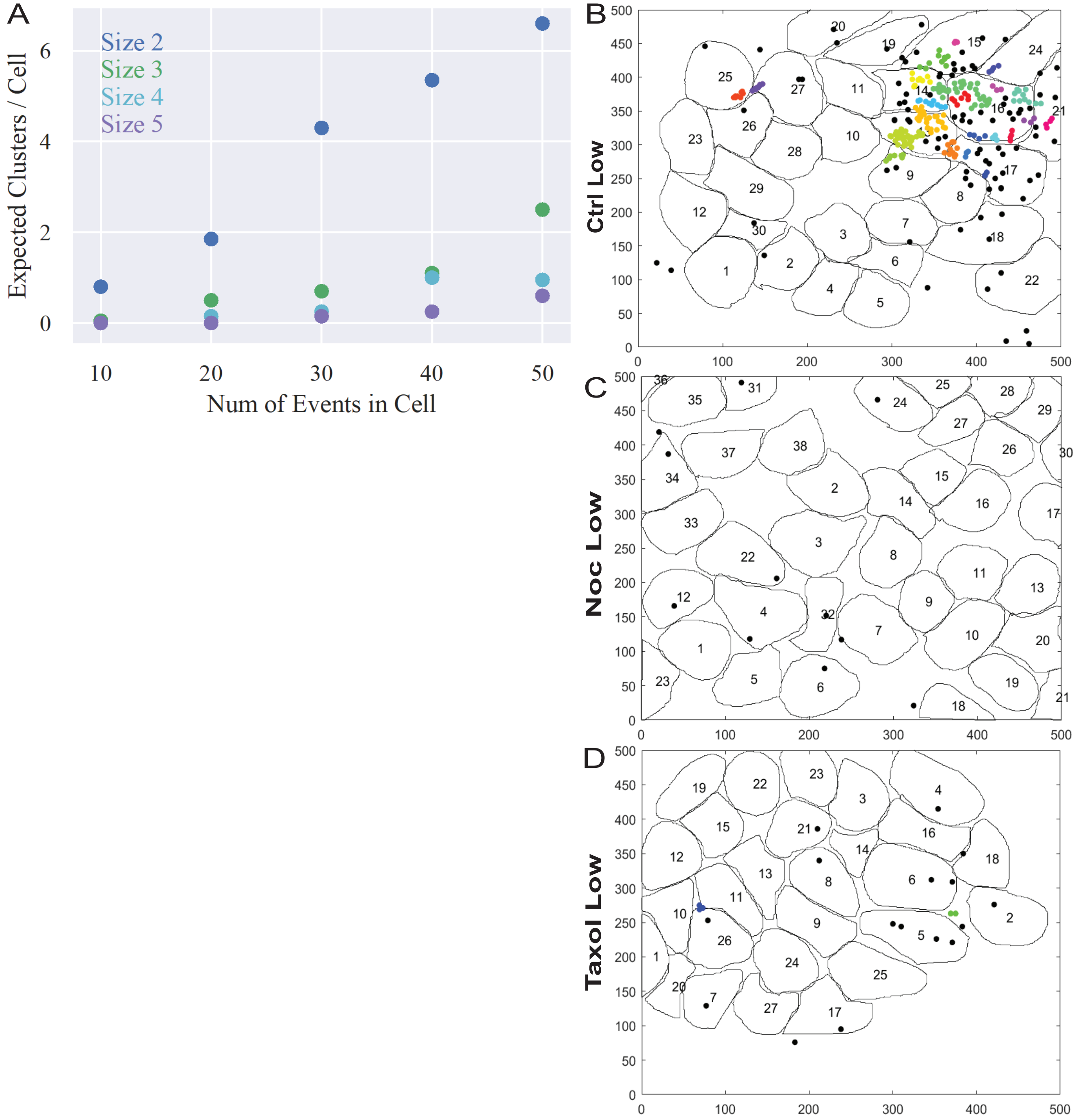

### Figure 16 figure supplement 1

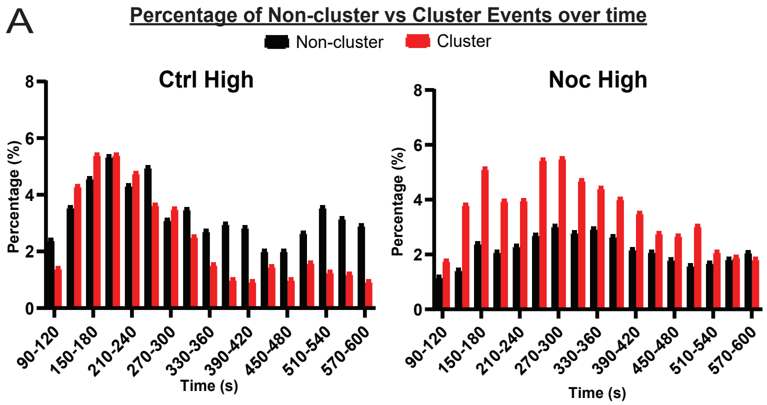
